# Per-Pathogen Virulence of HIV-1 subtypes A, C and D

**DOI:** 10.1101/2022.03.18.484874

**Authors:** Judith A Bouman, Colin M Venner, Courtney Walker, Eric J Arts, Roland R Regoes

## Abstract

HIV-1 subtypes differ, among other things, in their clinical manifestations and the speed in which they spread. In particular, the frequency of subtype C is increasing relative to subtype A and D. We aim to investigate whether HIV-1 subtype A, C and D differ in their per-pathogen virulence and to what extend this can explain the difference in spread between these subtypes.

We use data from the Hormonal Contraception and HIV-1 Genital Shedding and Disease Progression among Women with Primary HIV Infection (GS) Study. For each study participant, we determine the set-point viral load value, CD4^+^ T cell level after primary infection and CD4^+^ T cell decline. Based on both the CD4^+^ T cell count after primary infection and CD4^+^ T cell decline, we estimate the time until AIDS for each individual. We then obtain our newly introduced measure of virulence as the inverse of the estimated time until AIDS. This new measure of virulence has an improved correlation with the set-point viral load compared to the decline of CD4^+^ T cells alone. After fitting a model to the measured virulence and set-point viral load values, we tested if this relation varies per subtype. We found that subtype C has a significantly higher per-pathogen virulence than subtype A. Based on an evolutionary model, we then hypothesize that differences in the primary length of infection period cause the observed variation in the speed of spread of the subtypes.

**Author summary:** HIV-1 subtype C is currently spreading relatively fast in various parts of the world. Data from a study that followed many women infected with different HIV-1 subtypes (A, C and D) before they started treatment shows that neither their viral load nor the disease duration are increased for subtype C compared to subtype A and D. Thus, it seems that subtype C does not have a transmission advantage, neither per contact nor due to longer infection, making the observed relative rise in subtype C a puzzle. We used the same data to test if subtype C has optimized its potential to spread by decreasing the disease duration per unit of viral load (per-pathogen virulence) compared to subtype A and D. However, we find that subtype C has a significantly higher per-pathogen virulence than subtype A and D. This result makes the rise of subtype C even more counter-intuitive.

In a last step, we develop an evolutionary model, in which we synthesize all our empirical results. With this model we can show that the most likely explanation for the global spread of subtype C is a difference in the duration of primary infection between the subtypes.

## Introduction

HIV-1 is genetically classified into groups M (Major), O (Outlier), N (non-M, non-O) and P [1]. Group M is responsible for the global pandemic of HIV-1 and different subtypes have been defined within this group: A-D, F-H, J,K [1]. Subtype B is most common in the Americas and Europe, whereas subtypes A, C, D and a recombinant form (CRF02-AG) are the most prevalent on the African continent [2]. Over the past years, the relative frequency of subtype C has been increasing in many parts of the world, including Brazil [3]–[6], Kenya, Uganda and east Asia, even though the general incidence of HIV-1 has decreased [7].

The subtypes of HIV-1 differ in their clinical manifestations. Individuals infected with different subtypes vary in their rate of disease progression [8]–[13], treatment failure, and drug resistance evolution [14]. In particular, subtype D correlates with faster disease progression and higher treatment failure and drug resistance as compared to subtype A [9], [14], [15]. The rate of disease progression of subtype C compared to type A and D is still debated [8], [11], [13], [16].

Most studies reporting on differences in disease progression between subtypes rely on CD4^+^ T cell decline [8], [16], [17], which has been found to be an independent predictor of disease progression in HIV-1 cohort studies [18], [19]. The CD4^+^ T cell decline is defined as the number of CD4^+^ T cells depleted per year throughout the course of HIV-1 infection. Even though the time until disease or death is the most direct measure of virulence of any infection, the CD4^+^ T cell decline is commonly used as a surrogate measure [18], [19]. The reason thereof is threefold. First, measuring the time to disease or death is complicated because of uncertainties in determining the time of infection. Secondly, available treatment makes it unethical to study the natural course of an infection [20]. Thirdly, the rate of decline can be calculated on a much shorter time scale than the direct observation of disease progression requires.

In most infections, virulence is correlated with pathogen load [21], [22]. For an HIV infection, the viral load is measured as the approximately constant level it attains during chronic infection. This set-point viral load level correlates strongly with the time until disease [23]. The set-point viral load also correlates with the transmission probability per contact [24], [25] and is therefore subject to a trade-off between the transmission and the duration of the infection, i.e. the transmission-virulence trade-off [26].

To capture how virulent pathogens are irrespective of the viral load they attain, evolutionary ecologists developed the concept of per-pathogen pathogenicity [27]. Even though we use the concept of per-pathogen pathogenicity as introduced by Råberg and Stjernman, we prefer to refer to it as *per-pathogen virulence*. The reason is the consensus that pathogenicity is not a continuous measure, but rather the quality or state of being pathogenic, whereas virulence is [28]. There is qualitative evidence, supporting that HIV-1 subtypes differ in their per-pathogen virulence. Baeten et al. (2007) observed a faster disease progression for individuals infected with subtype D than for individuals infected with subtype A, even though they attained similar set-point viral load values [9]. The classical transmission-virulence trade-off predicts that a subtype with a high per-pathogen virulence has an evolutionary disadvantage of spreading compared to a subtype with a low per-pathogen virulence, because it has an additional cost in terms of virulence for increasing the viral load.

In this study, we use data from the Hormonal Contraception and HIV-1 Genital Shedding and Disease Progression among Women with Primary HIV Infection (GS) Study [8], [29]–[31] to investigate whether HIV-1 subtypes indeed differ in their per-pathogen virulence. In our cohort, we found that the first CD4^+^ T cell measure after primary infection differs by subtype and correlates with the CD4^+^ T cell decline. As a result, the expected time until AIDS depends not only on the CD4^+^ T cell decline, but also on the CD4^+^ T cell level at the end of primary infection. This forced us to abandon using the CD4^+^ T cell decline as the measure of virulence and to develop an alternative quantitative measure of HIV-1 virulence that incorporates the CD4^+^ T cell level immediately after primary infection. This alternative measure is more comparable to the direct observation of the time until death or AIDS than the decline of CD4^+^ T cells alone. Using this measure we can establish that HIV-1 subtype C has a significantly higher per-pathogen virulence than subtype A.

We put this conclusion into context with the recent geographical expansion of subtype C infections. This observation is contradicted by the prediction from the classical transmission-virulence trade-off that a subtype with a high per-pathogen virulence has an evolutionary disadvantage. To explain this contradiction, we expand the classical transmission-virulence framework by including the different transmission rates during primary and chronic infection. Based on the expanded framework, we conclude that the duration of primary infection could play a crucial role in the relative fast spread of subtype C.

## Methods and Materials

### Study Population

303 Participants from Uganda and Zimbabwe enrolled between 2001 and 2009 in the GS-study, after they got infected with HIV-1 during participation of the Hormonal Contraception and the Risk of HIV Acquisition (HC-HIV) Study [8], [29]–[32]. Consequently, all participants are woman of child-bearing age. Ethical approval was obtained from the Institutional Review boards (IRBs) from the Joint Clinical Research Centre and UNST in Uganda, from the University of Zimbabwe, from the University Hospitals of Cleveland, and recently, from Western University. Protocol numbers and documentation of these approvals/renewals are available upon request. Consent was not obtained as data were analyzed anonymously.

The participants were followed for an average of 5 years, during which CD4^+^ T cell and viral load levels were monitored. Following WHO recommendations at the time of the study, starting from 2003, women who had two consecutive CD4^+^ T cell counts below 200 cells/*µ*l or showed symptoms of AIDS were offered highly active anti-retroviral therapy [29].

Observations obtained during treatment are filtered out of the data, as they do not reflect the natural dynamics of the infection. Additionally, data-points observed during the primary infection are excluded, as the set-point viral load and the CD4^+^ T cell decline have to be determined during the chronic part of infection.

After filtering the data, we include participants for whom at least 3 viral load measurements and 4 CD4^+^ T cell counts are available. Also, CD4^+^ T cell counts have to span more than 180 days. An overview of all included participants, their country of origin and HIV-1 subtype is shown in Table 1.

**Table 1:**
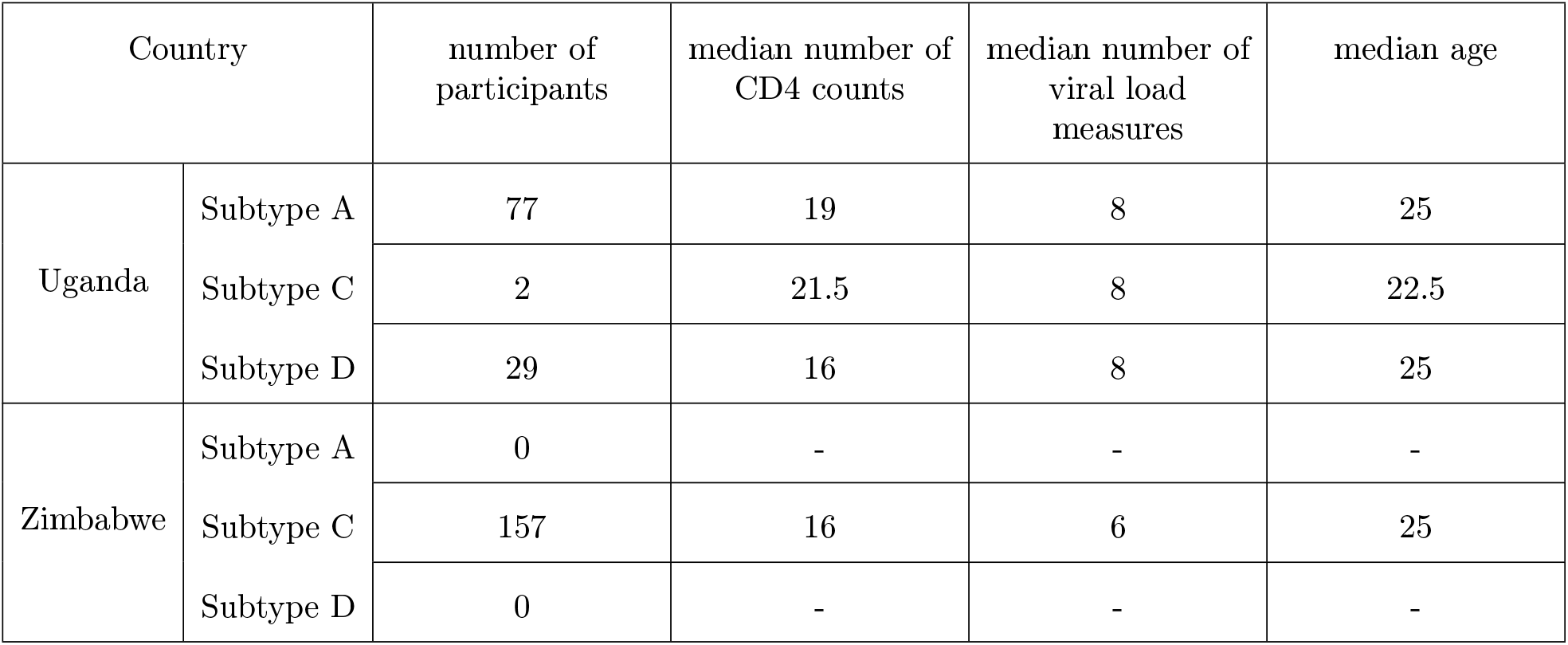
Overview of study participants.

| Country |  | number of participants | median number of CD4 counts | median number of viral load measures | median age |
| --- | --- | --- | --- | --- | --- |
| Uganda | Subtype A | 77 | 19 | 8 | 25 |
|  | Subtype C | 2 | 21.5 | 8 | 22.5 |
|  | Subtype D | 29 | 16 | 8 | 25 |
| Zimbabwe | Subtype A | 0 | - | - | - |
|  | Subtype C | 157 | 16 | 6 | 25 |
|  | Subtype D | 0 | - | - | - |

### Linear mixed-effects Model of CD4^+^ T cell Counts

The measurements of CD4^+^ T cells are used to calculate the CD4^+^ T cell decline for each participant. To reduce the impact of the within person variance and take advantage of patterns shared between participants, we used a linear mixed-effect model for the repeated CD4^+^ T cell measures with participant number as a random effect and time as a fixed effect. The decline of the CD4^+^ T cell counts and the CD4^+^ T cell count after primary infection are then obtained as the Best Linear Unbiased Predictors (BLUPs) of the slope and intercept of the fit for each individual [13].

### Rate of Disease Progression

We defined a new measure of the virulence that incorporates both the level of CD4^+^ T cells after primary infection (*CD*4^+^(0)) and the CD4^+^ T cell decline (Δ*CD*4^+^). We call this measure the “rate of disease progression”.

In a first step, we estimate the duration of the chronic infection in an infected individual (*t*_*I*_) by a linear extrapolation of the CD4^+^ T cell levels at the end of primary infection (*CD*4^+^(0)) to the AIDS-defining level of 200 cells per *µl* blood:

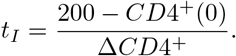

A visual intuition for this extrapolation is shown in Figure 1. The decline in CD4^+^ T cell is assumed to be linear, this assumption has been tested by comparing a linear fit to a two-phase linear, exponential and power-law fit.

**Figure 1:**
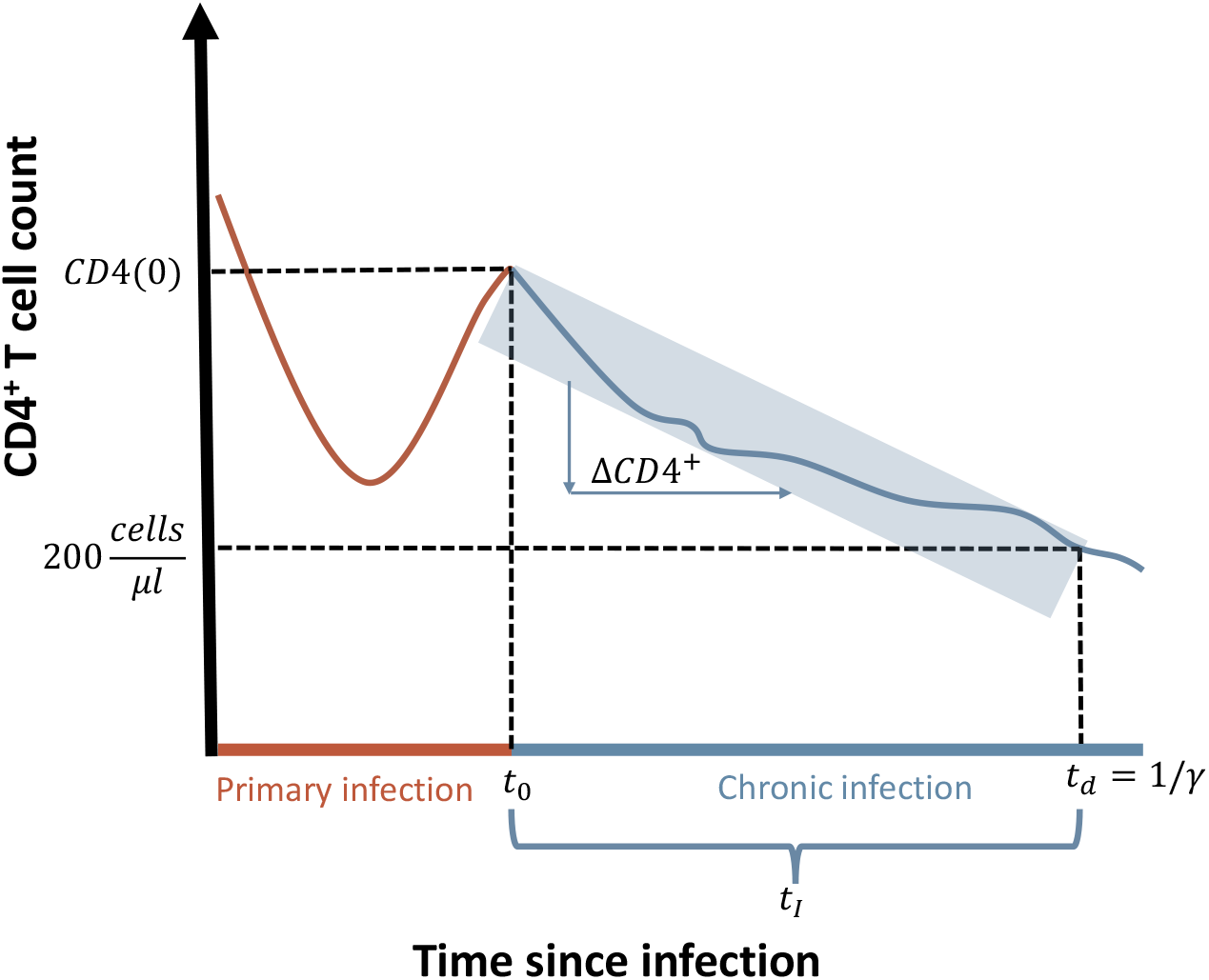
Schematic representation of the CD4^+^ T cell counts during an HIV-1 infection. Indicated are; *t*_0_, the end of primary infection; *CD*4(0), the CD4^+^ T cell level after primary infection; Δ*CD*4^+^, the decline of CD4^+^ T cells during chronic infection; *t*_*d*_, the time at which AIDS starts; and *γ*, the inverse of the time until AIDS.

To obtain the full time until AIDS (*t*_*d*_), we add the duration of primary infection to *t*_*I*_. Lastly, we calculate the rate of disease progression by taking the inverse of the extrapolated total time until AIDS.

### Per-Pathogen Virulence

To calculate the per-pathogen virulence, we need to correct the rate of disease progression for the viral load of each individual. Therefore, we first calculate the set-point viral load by taking the geometric mean of the viral load measurements for each participant. We then investigate the relationship between the rate of disease progression and the set-point viral load. The description of this relation is given in Equation 1. This relationship includes the age of the individual, as previous studies have shown that age influences the relation between virulence and set-point viral load for HIV-1 [33]. Moreover, it contains a coefficient for subtype C and a parameter for the difference with subtype A and D. These coefficients represent the per-pathogen virulence of the subtypes.

The exponent in Equation 1 is first determined without the other model parameters with the ’nls’ function in R [34]. We estimated the exponent to be 3.12 (95%*CI* : (2.35 − 3.96)). We then fixed this exponent to estimate the other model parameters with the ’lm’ function in R [34].

### Software

All data analyses and calculations have been done in R, version 1.1.456 [34], [35]. The linear mixed effect models are executed with the ’lmer’-function of the ’lmer’ package [36].

## Results

### CD4^+^ T cell Decline is linear

CD4^+^ T cell decline is a common way to measure the virulence of a HIV-1 infection. As a first step in our analysis, we investigated if the CD4^+^ T cell level declines linearly over time. To this end, we statistically compared models assuming a linear decline to alternative temporal relationships: two-phase linear, exponential and power-law. We found the highest statistical support for the linear decline model. This corroborates the assumptions of previous analyses [33], [37], [38]. For the subsequent analysis, we therefore assumed that the decline of CD4^+^ T cells is linear in time.

### CD4^+^ T cell Decline, CD4^+^ T cell level after Primary Infection and set-point Viral Load are Associated with HIV-1 Subtype

We are interested in subtype specific differences in the rate of disease progression and per-pathogen virulence. Before addressing these, we investigate potential subtype differences in the CD4^+^ T cell decline, set-point viral load, and CD4^+^ T cell level after primary infection. This is necessary because estimates of the rate of disease progression and per-pathogen virulence rely on these quantities.

Using the linear model for CD4^+^ T cell decline, we estimated the CD4^+^ T cell level after primary infection and the rate of CD4^+^ T cell decline in each study participant as the best linear unbiased predictors in a linear mixed-effects model (see Materials and Methods). These CD4^+^ T cell decline estimates differ by the viral subtype causing the infection (see Figure 2 A). Specifically, the CD4^+^ T cell decline has been found to be significantly lower for participants infected with subtype C than for participants infected with subtype D (Wilcoxon rank test on the individual decline estimates, p-value = 0.037, see Figure 2 A). This is consistent with previous work using a GEE approach to measure rates of CD4^+^ T cell decline on the GS-dataset, i.e. that subtype C infected participants had slower CD4^+^ T cell decline from a lower CD4^+^ T cell level following viral load set point during early disease [8].

**Figure 2:**
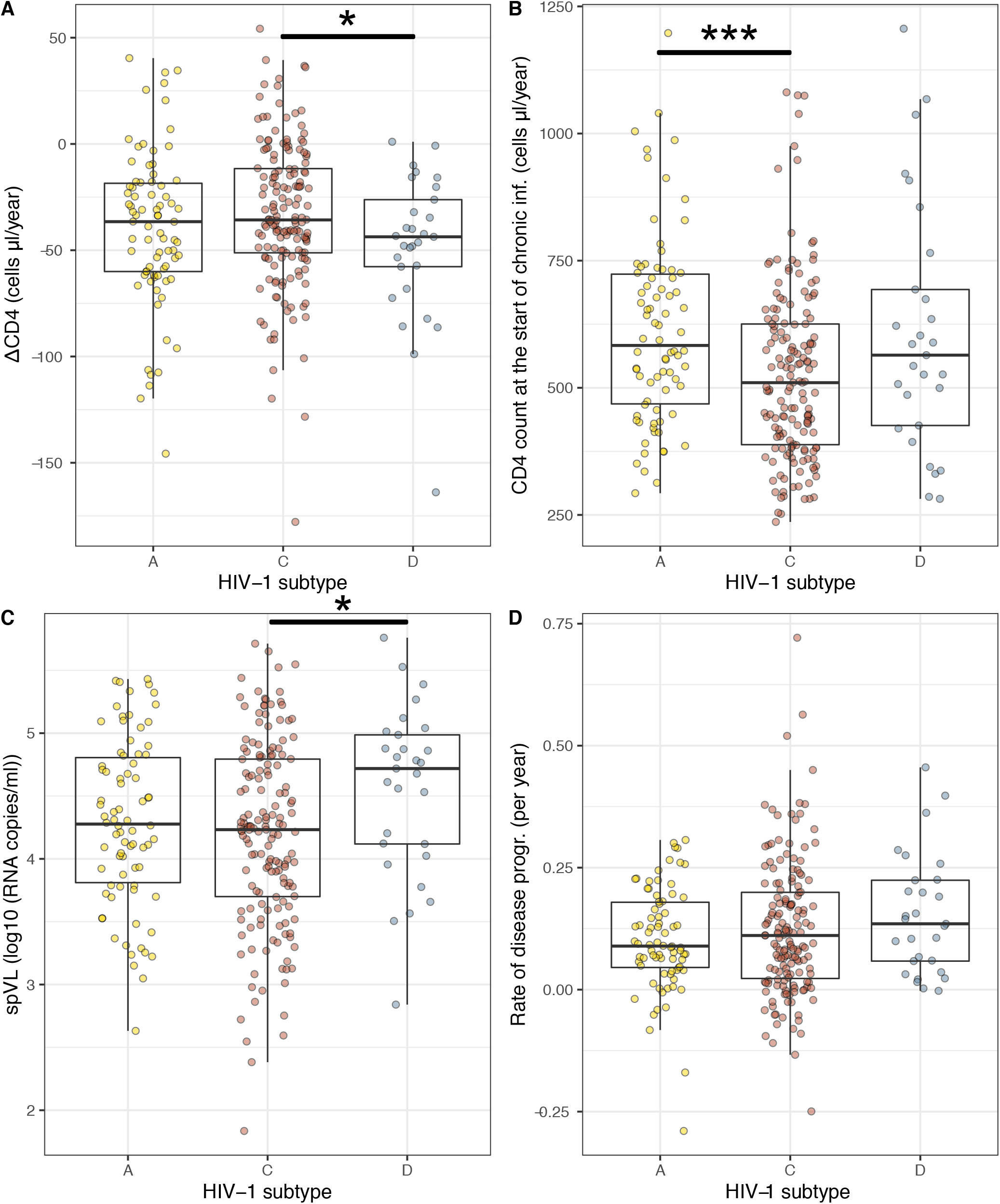
(A) CD4^+^ T cell declines per subtype, there is a significant difference between subtype C and D (p-value = 0.037). (B) CD4^+^ T cell counts at the start of the chronic infection per subtype, there is a significant difference between subtype A and C (p-value = 0.00056). (C) Set-point viral load per subtype, there is a significant difference between subtype C and D (p-value = 0.019). (D) Rate of disease progression per subtype, no significant differences are observed. 9

Also, we found that set-point viral loads vary per subtype, subtype C induces a significantly lower set-point viral load than subtype D (see Figure 2 C, Wilcoxon rank test, p-value = 0.019). This is in line with the difference in the CD4^+^ T cell decline, in the sense that the high set-point viral load of subytpe D compared to subtype A corresponds to a fast CD4^+^ T cell decline for subtype D.

However, not only the set-point viral load and CD4^+^ T cell decline differ by subtype, also the CD4^+^ T cell count after primary infection varies between these groups, see Figure 2 B. In this case we found a significantly lower value for subtype C than for subtype D (p-value = 0.00056). This has been observed before by Lovvorn et al (2016). Moreover, we found that low levels of CD4^+^ after primary infection correlate with low CD4^+^ T cell decline (Spearman Correlation = *−*0.14, p-value = 0.025).

### Per-Pathogen Virulence is higher for Subtype C than for Subtype A

The associations of HIV-1 subtypes with the CD4^+^ T cell level after primary infection and the CD4^+^ T cell decline have conflicting implications on the rate of disease progression. While the lower CD4^+^ T cell level after primary infection for subtype C will accelerate the progression toward disease, the slower CD4^+^ T cell decline will counteract this effect.

To incorporate these conflicting influences, we developed a measure of virulence — the “rate of disease progression” — that depends on both, the decline of CD4^+^ T cells and their level immediately after primary infection (see Material and Methods). In contrast to the CD4^+^ T cell decline, the rate of disease progression does not significantly vary by subtype (see Figure 2 D).

The subtype specific per-pathogen virulence can be quantified as the coefficient of the rate of disease progression against the set-point viral load (see Materials and Methods). Formally, this regression can be expressed as the following equation for the rate of disease progression, *γ*:

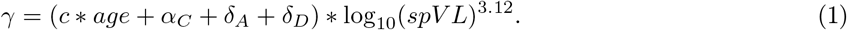

In this expression, the rate of disease progression is a function of the set-point viral load *spV L* and parameter *c* captures the age-dependence. The per-pathogen virulence is measured by parameters *α*_*C*_, *δ*_*A*_, and *δ*_*D*_. Hereby, *α*_*C*_ is the per-pathogen virulence of subtype C, and *δ*_*A*_ and *δ*_*D*_ denote the difference between the per-pathogen virulence of subtype A and D to subtype C, respectively. The best estimates for these parameters are listed in Table 2. The model explains 27.4% of the variance in the rate of disease progression.

**Table 2:** Parameters of the per-pathogen pathogenicity model.

| Parameter | Value | p-value |
| --- | --- | --- |
| $c$ | 3.340e-5 | 0.0246 |
| $\alpha_C$ | 1.326e-3 | <2e-16 |
| $\delta_A$ | -3.544e-4 | 0.0149 |
| $\delta_D$ | 2.507e-5 | 0.8888 |

We found that HIV-1 subtype C has a significantly higher per-pathogen virulence than subytpe A (p-value = 0.015). In practice, this will translate into an almost 50% increase of predicted time to disease for subtype A compared to subtype C for a set-point viral load of 3 *∗* 10^5^ RNA copies per ml.

### Evolutionary Implications

We have observed that subtype C has a significantly higher virulence than subtype A for similar set-point viral load values. This has implications for the competition of these subtypes on the population level.

Generally, it is assumed that viruses evolve to maximize their transmission potential — a combination of the transmission probability per contact and the length of the infectious period [39]. In HIV, the set-point viral load is strongly correlated to the transmission probability [26] and inversely correlated to the duration of the infectious period [40], [41]. Hence, there is a trade-off between transmission and virulence [26]. The per-pathogen virulence influences the shape of this trade-off. Subtype C, which has a higher per-pathogen virulence than subtype A, endures a higher cost in terms of virulence for increasing the transmission with a higher viral load (see Figure 4). Because the set-point viral load is similar for the two subtypes, we expect the transmission for subytpe A to be higher than for subtype C.

**Figure 3:**
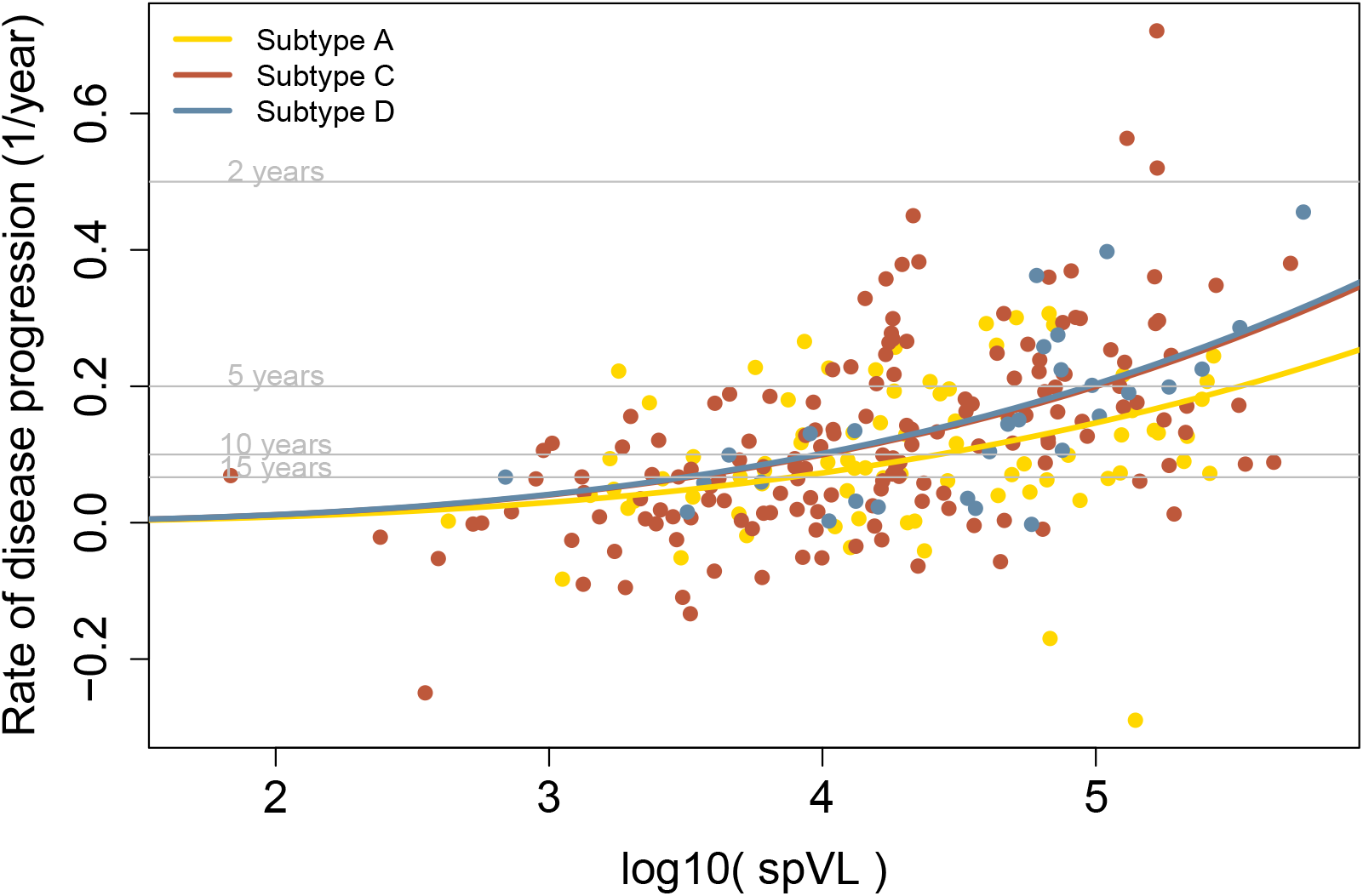
Relation between rate of disease progression and 10 base logarithm of the set-point viral load.

**Figure 4:**
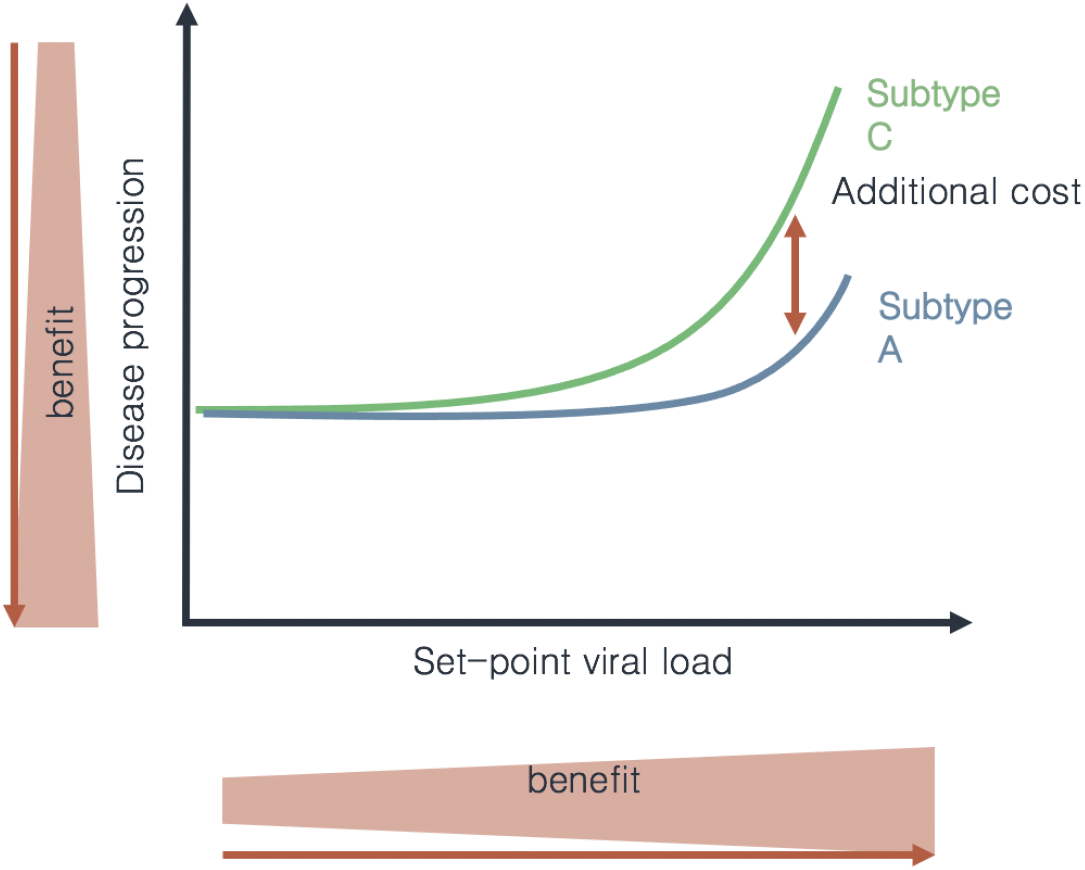
The relation between the set-point viral load and the disease progression is shown for subtype A and C. A high set-point viral load is beneficial for the transmission potential, whereas a high rate of disease progression is disadvantageous. The higher per-pathogen virulence of subtype C leads to an additional cost in terms of disease progression for an increase of the set-point viral load.

This expectation, however, is not consistent with the increase of the relative frequency of subtype C currently observed in different parts of the world [4], [5], [7], [42]. The classical transmission-virulence trade-off that gave rise to the expectation assumes a constant transmission rate during the course of the infection. However, for HIV-1, it is known that the transmission rate is significantly higher during primary infection than during chronic infection [43]. Both the primary and the chronic part of the infection account for around half of the transmission potential, even though the chronic part of infection takes much longer (see Figure 5 A). To take this into account, we extended the classical transmission-virulence trade-off by explicitly including the transmission potential of the primary and chronic part of the infection. As a result, this extended model contains four parameters: the transmission rate and duration of primary infection and the transmission rate and duration of chronic infection.

**Figure 5:**
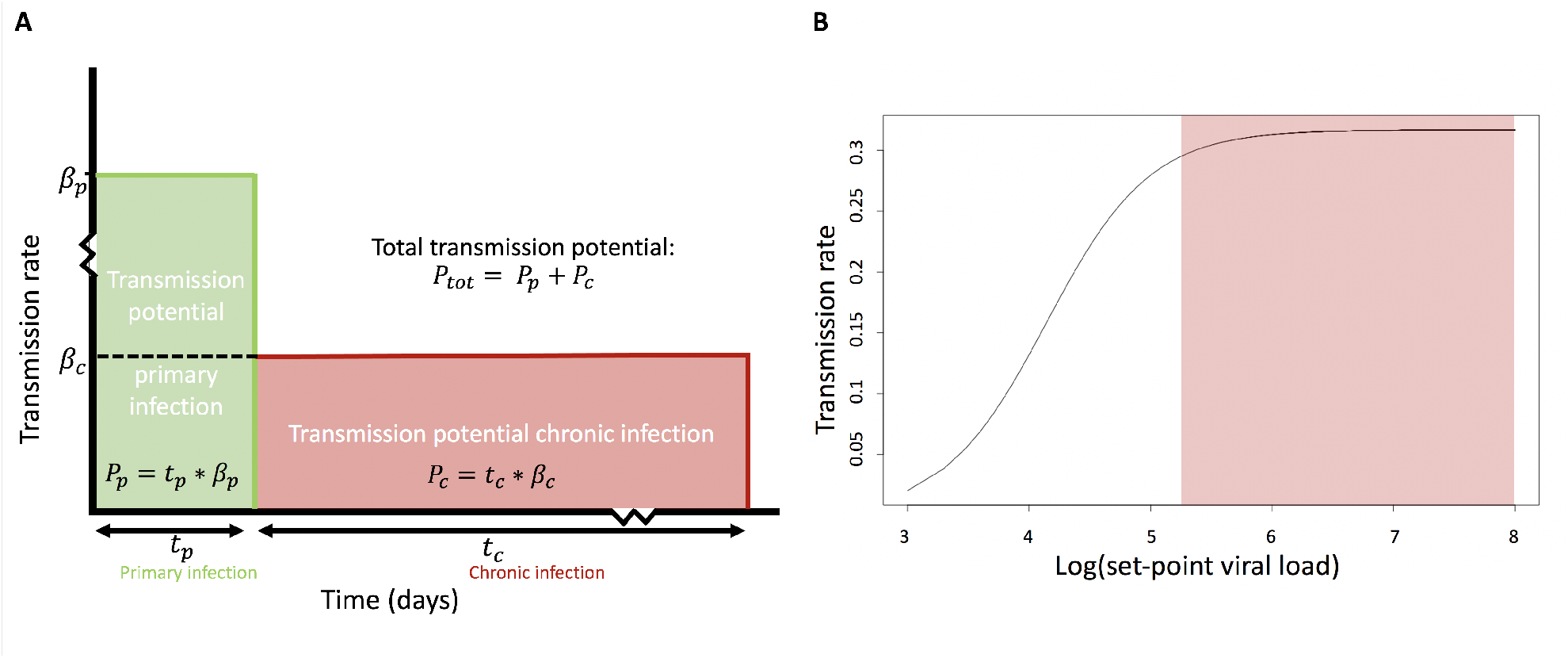
Panel A is adapted from Hollingsworth et al. (2008) and shows the different parameters that contribute to the total transmission potential: the duration of primary infection, the transmission rate during primary infection, the duration of chronic infection and the transmission rate during chronic infection. Panel B is adapted from Fraser et al. (2007) and shows the relation between the set-point viral load and the transmission rate.

We first consider the chronic part of the infection. The GS-dataset shows no difference in duration of chronic infection based on our measure of virulence. Moreover, we observed no significant difference in set-point viral load between subtype A and C and therefore no difference in transmission probability per contact is expected. From these two observations, we conclude that the total transmission potential during chronic infection is comparable between subtype A and C. As a result, the difference in spread between the two subtypes is most likely explained by a difference in the transmission potential during primary infection.

We then evaluate the transmission rate during primary infection using the relation between the viral load and the transmission rate determined by Fraser et al (see Figure 5 B) [26]. For simplicity, we assume the same relation to be valid during primary infection. Robb et al. showed that the peak viral load during primary infection has a mean value of 6.7 log_10_ *copies/mL* (range, 4.5 to 8.5) [44]. For viral load values this high, the transmission rate has saturated and is unlikely to vary, even when the peak viral load varies substantially (Figure 5). Additionally, Campbell et al. argue that viral loads are similar during primary infection for subtype C and non-C HIV-1 infections [45]. Thus, it is very unlikely that the difference in spread of subtype C and A is caused by a difference in transmission rate during primary infection that is driven by difference in peak viral load.

The duration of the primary infection is the only parameter that we cannot show to be similar between the studied subtypes based on available data. A relatively small difference in duration of primary infection could already lead to a significant increase of the transmission potential of subtype C compared to subtype A, as primary infection is responsible for around half of the total number of transmissions. Thus, based on this evolutionary consideration we conclude that the reason for the increase of the relative frequency of subtype C is possibly a longer duration of primary infection. As a consequence, the higher per-pathogen virulence of subtype C could be adaptive because it extends the primary infection period.

## Discussion

Our study was motivated by the observation that the frequency of subtype C increases relative to subtype A and D in many regions of the world [3]–[7]. There are two possible explanations for this observation: either subtype C has a transmission advantage and its relative increase is the result of selection, or the increase is due to stochastic fluctuations in prevalence [46]. In our study, we investigate if there is evidence for a transmission advantage of subtype C that could add support to the selection hypothesis.

We found that HIV-1 subtype C, compared to subtype A, has a higher per-pathogen virulence, a term which we prefer over the more commonly used per-pathogen virulence. This has been established on the basis of a newly defined measure of virulence — the rate of disease progression — which combines both the decline of CD4^+^ T cells and the CD4^+^ T cell level after primary infection. We had to incorporate this measure of virulence, because CD4^+^ T cell levels after primary infection varied significantly between subtypes and this counteracted the disease predictions based solely on the decline of CD4^+^ T cells.

The higher per-pathogen virulence of subtype C does not provide any selective advantage that could explain its increase in frequency. Rather, it constitutes a selective disadvantage. To explore if, despite a higher per-pathogen virulence, subtype C could have an overall transmission advantage, we extended the classical transmission-virulence trade-off model by the different contribution of primary and chronic infection to transmission. According to this evolutionary consideration, the only conceivable way to maintain a selective advantage of subtype C is a longer duration of primary infection. At this stage, we are not aware of any empirical evidence for a longer primary infection of subtype C. Neither the clinical data we used in the present work nor previous studies did provide any information on this aspect. Thus, the longer duration of primary subtype C infection is a hypothesis awaiting future testing.

The subtype specific differences in CD4^+^ T cell level after primary infection which we found are also observed between the countries of origin [13], [47] (supplementary Figure S1). Thus, the differences in CD4^+^ T cell level after primary infection could be caused by a geographic variation in CD4^+^ T cell level before seroconversion rather than by differential CD4^+^ T cell depletion by each subtype during primary infection. Lovvorn et al. reported this for the GS-data in 2010 [47]. They also tested for differences in CD4^+^ T cell level in HIV negative subjects from Zimbabwe and Uganda. Using a Mann-Whitney-Wilcoxon test, they report a country specific difference in the CD4^+^ T cell level after primary infection [47]. However, this difference in CD4^+^ T cells is larger for individuals infected with HIV-1 than for those that are HIV-naive [47]. Another cohort study testing for baseline CD4^+^ T cell differences between African countries is described by Karita et al. [48]. They record no significant differences between the countries in their cohort. Thus, the origin of the subtype specific difference in the CD4^+^ T cell level after primary infection is not clear yet.

As a result of the counteracting effects of the CD4^+^ T cell level after primary infection and the CD4^+^ T cell decline on the predicted time of disease, we did not find an association of the subtype on the rate of disease progression, while we do find an association of subtype on the CD4^+^ T cell decline. Several other studies report subtype associations with the time until disease or death [9]–[12], [14]–[16], [49], [50]. In particular, these studies found that individuals infected with subtype D progress to AIDS faster than individuals infected with subtype A [9], [12], [14]–[16], [49], [50]. In our analysis, we also find a trend that subtype D infected individuals progress to AIDS faster than individuals infected with subtype A, however this difference is not significant. This could be due to the relatively small number of individuals carrying subtype D in our cohort.

A comparison of subtype A and D with C is less studied, as data which contain all three types are rare. The studies that do combine subtypes A, C and D are ambiguous in their conclusion. Some find no difference in disease progression for subtype C compared to A and D when considering direct observations of the time until death or AIDS or the CD4^+^ T cell decline corrected for baseline CD4^+^ levels [11], [16]. However, based on the CD4^+^ T cell decline alone, Amornkul et al (2013) find a faster rate of disease progression for subtype C compared to A and Venner et al (2016) find a slower rate of disease progression for subtype C compared to A and D [8], [13], [46]. Using the same clinical data as Venner et al (2016), we confirm their finding based on CD4^+^ T cell decline. Additionally, we extend it with the result that our more considerate measure of virulence – the rate of disease progression – does not vary between the studied subtypes, in accordance with the observations based on the direct observation of time until death or AIDS [11]. We perceive this agreement in results as an indication that the rate of disease progression is a more meaningful measurement of virulence than the CD4^+^ T cell decline alone.

There is a also a conceptual merit for the use of our new measure of disease progression: if the slow CD4^+^ T cell decline during chronic infection start at a higher level it will take longer to reach the AIDS defining level of 200 CD4^+^ T cell per ml blood. Any reasonable measure of virulence therefore needs to take the level of CD4^+^ T cells after primary infection into account if there is variation in this level. In a cohort in which there is no such variation, the CD4^+^ T cell decline might still be a valid surrogate of disease progression.

The validity of our new virulence measure is further supported by the finding that the set-point viral load explains more of the variance in the rate of disease progression compared to CD4^+^ T cell decline (*R*^2^ = 0.27 compared to *R*^2^ = 0.1191). Our low *R*^2^ value for the relation between the set-point viral load and the decline of CD4^+^ T cells is consistent with multiple other cohort studies [33], [42]. On the other hand, high correlations have been found for the relation between set-point viral load and time until AIDS and death [23]. The increase in explained variance for the rate of disease progression makes us believe that including the CD4^+^ T cell count after primary infection results in a measure of virulence that is more predictive of the observed time until disease.

To investigate the per-pathogen virulence of HIV-1 subtype A, C and D, we combined the virulence and set-point viral load measurements. This led to the conclusion that even though disease progression does not differ per subtype, the disease progression for similar set-point viral load values is faster for subtype C compared to subtype A. The main driver of this difference are individuals infected with subtype C that have very low CD4^+^ T cell values after primary infections, leading to fast disease progression. This finding is in agreement with the observation of Baeten et al. (2007) [9], who found higher survival probabilities for individuals infected with subtype A than for individuals infected with subtype D, for similar set-point viral loads.

The evolutionary framework we applied to our observations included both the transmission potential of the primary and chronic part of the infection and allowed us to hypothesize that the faster epidemiological spread of subtype C compared to the other subtypes is most likely caused by differences in the length of primary infection. To obtain the hypothesis, we assumed that the relation between the viral load within the host and the transmission rate is the same during primary infection as it is for the chronic part of the infection. However, it is unlikely that the higher transmission during primary infection is solely caused by a higher amount of viral load inside of the host [51]. Many factors could influence this relation, such as the presence of viruses in compartments more relevant for sexual transmission of the disease. We cannot fully rule out the possibility that there are also subtype specific differences in the way in which viral load relates to the transmission rate during primary infection. However, currently there is no evidence for such an effect. Additionally, the observation of the subtype specific differences in CD4^+^ T cell level after primary infection is an indication that primary infection dynamics differ per subtype. Overall, we hypothesize that subtype specific differences during primary infection are the main driver of the distinguishable speed of spread of subtype C compared to subtype A.

## Ethical statement

Ethical approval of the GS-study was obtained from the Institutional Review boards (IRBs) from the Joint Clinical Research Centre and UNST in Uganda, from University of Zimbabwe, from the University Hospitals of Cleveland, and recently, from Western University. Protocol numbers and documentation of these approvals/renewals are available upon request [8].

## Acknowledgements

We would like to thank Art Poon and two anonymous reviewers for providing helpful feedback.

## Supplementary Material

**Figure S1:**
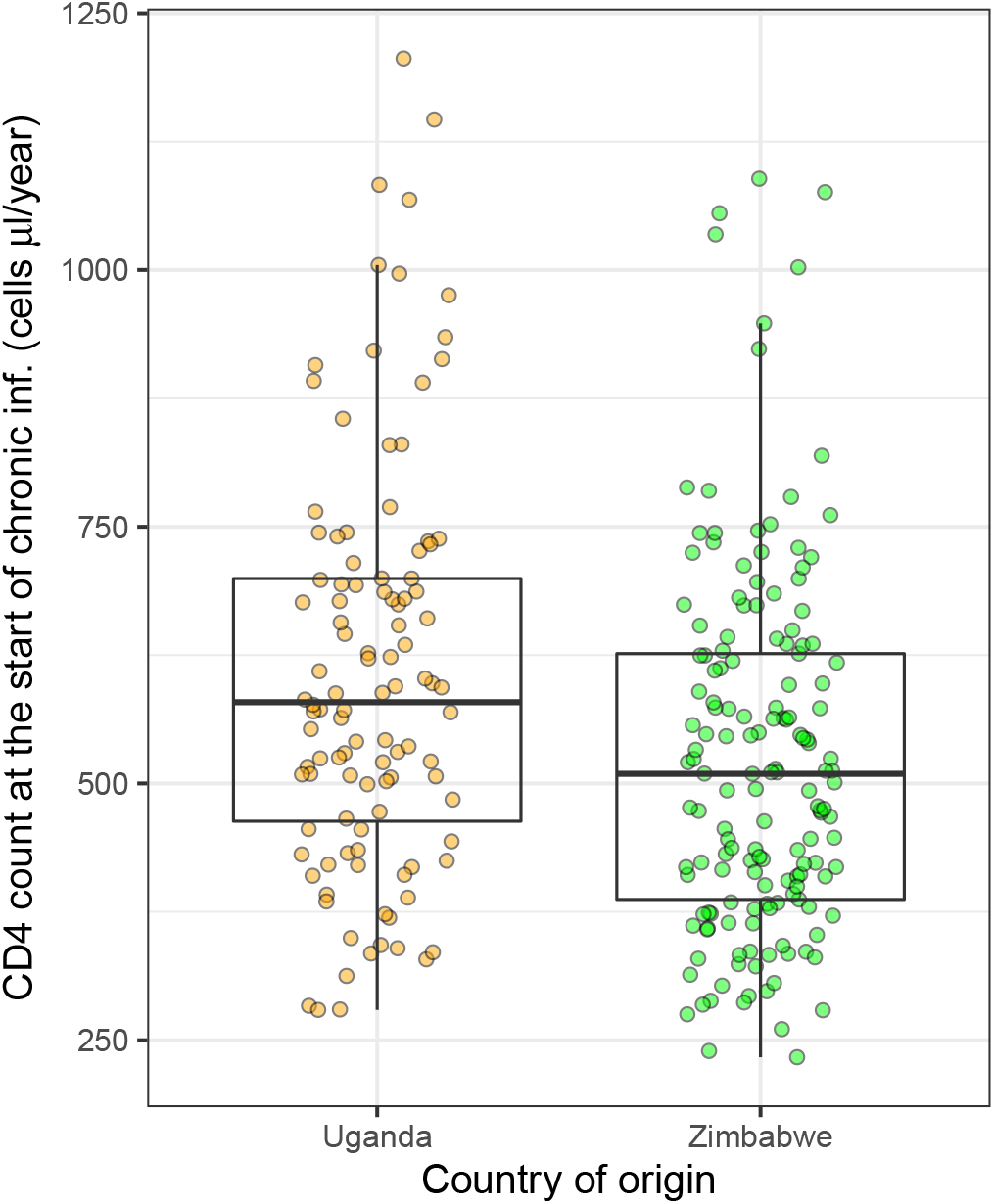
CD4^+^ T cell counts at the start of the chronic infection per country of origin, there is a significant difference between both countries (p-value = 0.004).

